# Cryptic haplotypes of “*Candidatus* Liberibacter africanus”

**DOI:** 10.1101/016410

**Authors:** Warrick R Nelson, Sandrine Eveillard, Marie-Pierre Dubrana, Joseph M Bové

## Abstract

“*Candidatus* Liberibacter africanus” (Laf) has long been recognised as a causal agent of the devastating citrus disease huanglongbing (HLB) or citrus greening. This species is currently restricted to Africa, the Arabian Peninsula and some Indian Ocean islands and vectored by the African citrus psyllid, *Trioza erytreae*. Blotchy mottle on citrus leaves is characteristic of the disease.

Somewhat similar symptoms in the Rutaceous tree *Calodendrum capensis* (Cape Chestnut) resulted in the discovery of Laf outside commercial citrus crops in South Africa. This was classed as a subspecies of Laf (capensis, hence LafC). In subsequent surveys of both commercial citrus crops and *Calodendrum*, both natural and ornamental specimens, LafC was not found in the citrus crop, nor has Laf been found in *C. capensis*. HLB was reported from Madagascar in 1968 but no sequences from this source have so far been published.

Until fairly recently, only the reference 16S rRNA gene sequences of Laf (L22533) and LafC (AF137368) had been deposited in GenBank. Both of these reference sequences contain a number of unresolved nucleotides. Resolving these nucleotide positions by aligning against more recently available sequences, it becomes evident that these unresolved positions represent one percentage point difference in similarity between Laf and LafC. The originally reported 97.4% similarity is therefore incorrect based on this new information. Recalculating the similarity on the full length 16S rDNA sequence results in 99.54% similarity, a value too high to justify a subspecies status. LafC should therefore be reduced to that of a haplotype of Laf.

Further, the six 16S rRNA gene sequences currently available in GenBank identified as the species Laf separate into 2 haplotype groups. The 3 haplotypes of Laf are therefore LafA designated as the first accession sequenced (L22533), LafC for the former capensis subspecies and to recognise the prior use of this term, and LafB for the third haplotype not previously recognised. Thus the cryptic presence of 3 haplotypes is revealed by this review of the Laf 16S rDNA sequences.

## Introduction

Huanglongbing (HLB), also known as citrus greening, is a serious disease threatening the economic existence of many commercial citrus orchards. It has recently spread to the major production regions of Brazil, Florida and California (Wang & Trivedi, 2013). Currently, 3 species of “*Candidatus* Liberibacter”, named for the continent of first discovery, are associated with symptomatic commercial citrus plants (Bové, 2006), “*Candidatus* Liberibacter asiaticus” (Las) being the most geographically widespread (Bové, 2014).

“*Candidatus* Liberibacter africanus” (Laf) is associated with HLB throughout East Africa, the Arabian Peninsula and Indian Ocean islands (Bové, 2006; Bové, 2014), including Madagascar (Bové & Cassin, 1968). The Rutaceous tree, *Calodendrum capense* (Cape Chestnut) has also been discovered to be infected by “*Ca*. Liberibacter africanus subsp. capensis” (LafC)(Garnier *et al.*, 2000). This has subsequently been found to be widespread in South Africa where this tree is both native and grown ornamentally (Phahladira *et al.*, 2012). In spite of the subspecies being widespread, only Laf is present in symptomatic commercial citrus orchards in South Africa (Pietersen *et al.*, 2010). Other native Rutaceae species in South Africa also tested positive for Laf and LafC and possibly another variant of the species, but interpreted as a series of subspecies of Laf (Roberts *et al.*, 2014).

The original description of LafC was based on three gene fragments (Garnier *et al.*, 2000), of which the 16S rRNA gene is taxonomically the most important (Weisburg *et al.*, 1991; Woese & Fox, 1977). This 16S sequence (GenBank accession AF137368) was noted as having 97.4% similarity with Laf (L22533). Three percent divergence over the full gene sequence is a commonly recognised requirement for bacterial species designation (Stackebrandt & Goebel, 1994), thus the subspecies status for LafC. Closer inspection of these sequences revealed 6 unresolved nucleotides on LafC and 5 in the Laf reference sequences. There is thus potentially one percentage point of difference derived simply from these unresolved nucleotide positions. A review of this subspecies status is now feasible with the availability of more recent accession sequences.

## Methods and Materials

A literature review revealed further examples of both Laf and LafC 16S rDNA sequences, as well as some sequences for 16S-23S ISR, 23S and 50S rRNA gene regions. The megablast protocol in GenBank was used to find further closely related sequences.

These sequences were downloaded from the NCBI database (GenBank) and aligned in ClustalX 2.1 (Larkin *et al.*, 2007). Sequences for 16S rDNA from the genomes of two other Liberibacter species (“*Ca*. L. asiaticus” and “*Ca*. L. solanacearum”, Las and Lso respectively) were also downloaded for additional comparison. These were chosen as phylogenies generally place them closest to Laf (Nelson *et al.*, 2013).

DNA material stored from symptomatic leaves of sweet orange collected in Madagascar during 2011, previously reported to be Laf by PCR (Bové, 2014), was retested using the primer pair OA1/Oi2c (Jagouiex *et al.*, 1996) and the resultant 16S rDNA samples sequenced. The rpIJ component of the 50S rRNA gene was also sequenced.

## Results

On the 16S rRNA gene, 15 sequences identified as Laf or LafC were found. Three of these proved to contain only a very short component of the 16S gene and were discarded. Aligning the remaining 12 sequences showed a number of apparently random SNPs scattered across the accessions, presumably arising as sequencing errors, but the LafC sequences separated clearly.

In the sequence L22533 (Laf), there are 5 nucleotide positions labelled “N”. From within the aligned set of sequences, these can be unambiguously resolved to the nucleotides T, C, G, A and G. Similarly, in the sequence AF137368 (LafC), these also resolve unambiguously to A, A, A, G, C, C.

Laf sequences also aligned into 2 groups over 7 SNPs (Table 1), here designated as haplotypes LafA and LAfB. Excluding the occasional single sequence SNPs, the number of SNPs separating the 2 haplotypes and subspecies is outlined in Table 2. Although the sequence of each accession varied in length, these SNPs occur over an approximately 1100 bp length. Aligning the Laf sequences against the full 16S gene sequences for Las and Lso indicated that all the SNPs in Table 1 occur within the overlapping regions. Liberibacter 16S genes are approximately 1500 bp (Nelson *et al.*, 2015). Until full Laf sequences are available, the similarities reported here might prove to be high if further SNPs exist in the currently unknown regions. There are 8 SNPs between Las and Lso in these end regions of the gene, representing 0.53% difference. Assuming a similar number of SNPs between the Laf sequences in these end regions, we can calculate a likely worst-case scenario for the number of SNPs between the proposed haplotypes and the percentage similarity, assuming a 16S rRNA gene length of 1500 bp.

**Table 1.**
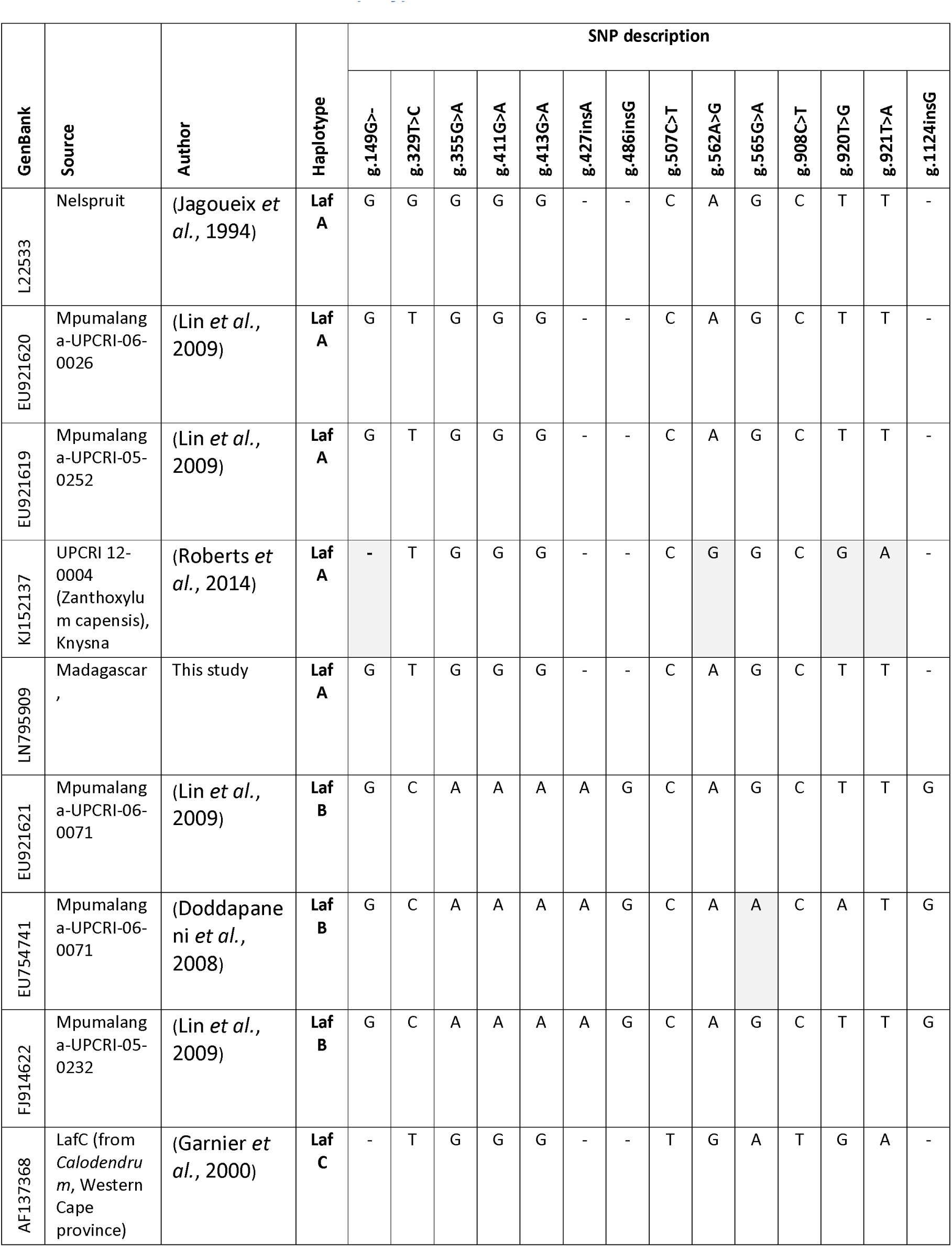
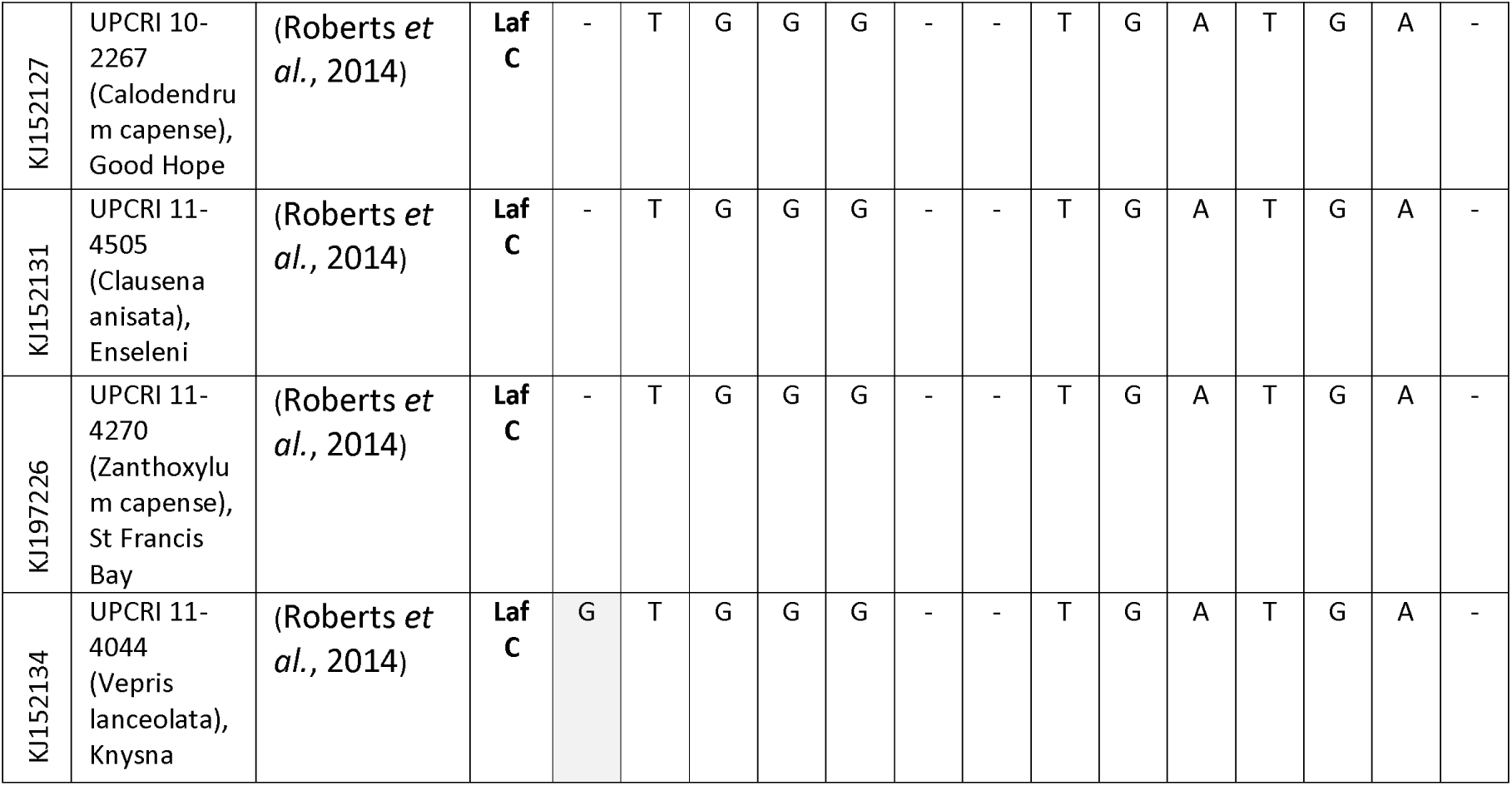
Proposed haplotypes and SNP differences on the 16S rRNA gene within “*Candidatus* Liberibacter africanus” (Laf) sequences. The nucleotide numbers count from the beginning of the reference sequence L22533 although the alignment was done using a 1057 bp length common to all sequences. Shaded SNPs are positions of ambiguity between the sequences within a series where the SNP is in the same position as the nucleotide in one of the other haplotypes.

**Table 2.** Percent identity similarity between “*Candidatus* Liberibacter africanus” (Laf) haplotypes based on 16S rRNA gene sequences, % similarity calculated on a 1500 bp length. An alternate % similarity figure is indicated in parentheses assuming a further 8 SNPs exists in the currently unknown segments of the gene.

| Comparison identity | SNPs | % similarity |
| --- | --- | --- |
| LafA:LafB | 7 (15) | 99.53 (99.0) |
| LafA:LafC | 7 (15) | 99.53 (99.0) |
| LafB:LafC | 14 (22) | 99.07 (98.54) |

Only 4 sequences across the 16S-23S ISR and 23S partial gene regions were found in GenBank (LAU61360, EU754741, FJ914622, JF819884, JF819885) for Laf and none for LafC. The LafC sequence has been mentioned as being closer to Las than Laf, because only one tRNA (alanine) is present in the ISR of Laf, while two tRNAs (alanine and isoleucine) occur in the ISR of Las and LafC (Garnier *et al.*, 2000).

Over a common segment of 479 bp on the 50S gene region (β-operon) there are 68 SNPs between Laf and LafC (14 and 4 accessions respectively)(Phahladira *et al.*, 2012; Pietersen *et al.*, 2010). The available sequence material in the 50S gene confirms the basic Laf/LafC separation, but we feel that separating LafA and LafB is as yet inconclusive with only 2 SNPs across the 4 accessions.

The 16S sequences derived from Madagascar material aligned with LafA and the rplJ sequence (GenBank) aligned with EF122255, a sequence with metadata indicating it is derived from commercial citrus material in South Africa.

## Discussion

The unresolved nucleotides in the sequences for Laf and LafC (11 positions in total) are readily resolved to their correct nucleotide by comparison against the other 10 sequences now available in GenBank. These newly resolved positions represent one percentage point of similarity between Laf and LafC, thus the originally described 97.4% (Garnier *et al.*, 2000) similarity becomes 98.4%. However, this similarity calculation was conducted on the known sequence proportion of the gene since at the time the full length of the gene was not known. Although the full 16S gene sequence for Laf is still not available, it is likely to be approximately 1500 bp in length as found in 4 other species of Liberibacter (Nelson *et al.*, 2015). Across the sequences currently available, only approximately 40 bases at the beginning and 30 at the end of the gene sequence remain unknown, thus allowing a closer estimate of similarity between the species and the subspecies (Table 2).

Variation across the 16S rRNA gene appears to be quite large with 14 SNPs apparent across the Laf/LafC accessions available, 7 of them between LafA/LafB and a different 7 between LafA/LafC, with all 14 being different between LafB/LafC. For comparison, Las appears to be homogenous across accessions on the 16S rRNA gene (Moreno-Enríquez *et al.*, 2014; Nelson, 2012), while for Lso 5 haplotypes have been described, with only 7 SNPs defining them on the 16S rRNA gene (Nelson *et al.*, 2011; Nelson *et al.*, 2013; Teresani *et al.*, 2014).

The unexpected separation into 2 groups on the 16S rRNA gene within the currently known Laf species suggests 2 haplotypes, here designated LafA and LafB (Table 2). No prior study has indicated that there were genetic differences within Laf on commercial citrus. This suggests 3 haplotypes within the Laf species, 2 primarily indicated by sequences derived from commercial citrus crops, and the previous subspecies designation “capensis” can be resolved to a haplotype but retaining the short form of LafC. Caution is required in this conclusion since the LafB designation is based on only 3 available sequences and 2 of these, although sequenced by different laboratories, are clearly derived from the same field sample. The metadata associated with these sequences give little indication of geographical or plant/insect host separation. For LafA/LafB they are primarily from commercial citrus crops and LafC from native Rutaceous species. This provides a stark contrast to the analogous situation in Lso where the haplotypes also express partial plant/insect host and geographic separation (Nelson *et al.*, 2011; Nelson *et al.*, 2013; Teresani *et al.*, 2014). If the A/B haplotypes can be confirmed, they indicate 2 separate pathosystem events from the respective currently unknown African native plant hosts into commercial citrus, since *Citrus* is not native to Africa (Beattie *et al.*, 2008), and LafC has yet to make this jump as it is not yet known from citrus (Phahladira *et al.*, 2012).

The 50S rRNA gene sequences show clear separation between Laf and LafC as expected, with only a suggestion on 4 of 15 Laf sequences showing SNPs. Caution is suggested in considering this indicative of LafA/B haplotypes as these SNPs occur together and only 4 bases from the end of the sequence. Further, there is no indication from the metadata to tie these 4 sequences together with the 3 suggested as LafB from the 16S gene.

Although previously known that Laf was the species responsible for HLB on Madagascar by both epidemiological and PCR studies (Bové & Cassin, 1968; Bové, 2014), this study confirms not only the species as Laf via both 16S and rplJ genes, but that it is also haplotype LafA. This very strongly suggests an incursion event of LafA from Africa to Madagascar, rather than a Gondwanan origin, in spite of the position of Madagascar in Gondwana between East Africa and India.

The very recent suggestion of a further three subspecies of Laf (Roberts *et al.*, 2014) can be resolved within this proposal of haplotypes rather than subspecies within Laf by giving them a biotype designation, recognising the current host plant differences. Laf subspecies vepridis is a biotype of LafA, while Laf subspecies zanthoxyli and Laf subspecies clausenae are biotypes of LafC.

## Conclusion

Instead of the species Laf and subspecies capensis, re-analysis of the phylogenetically important 16S rRNA gene suggests the subspecies is not that far removed from the species and should be revised to haplotype status. Further, existing sequences of Laf indicate that this species also comprises two haplotypes. Therefore two haplotypes (LafA and LafB) are known symptomatically in commercial citrus orchards while the third (LafC) is known only from very mild symptoms in native Rutaceous plants but not (yet) in citrus. Thus the cryptic presence of three haplotypes is revealed by this review of the Laf 16S rDNA sequences.

## References

Beattie, G., Holford, P., Mabberley, D., Haigh, A. & Broadbent, P. (2008). On the origins of *Citrus*, Huanglongbing, *Diaphorina citri* and *Trioza erytreae*. International Research Conference on Huanglongbing, Orlando, Florida, 25–57.

Bové, J. M. (2006). Huanglongbing: a destructive, newly-emerging, century-old disease of citrus. Journal of Plant Pathology 88, 7–37.

Bové, J. M. (2014). Heat-tolerant Asian HLB meets Heat-sensitive African HLB on the Arabian Peninsula. Why?. Journal of Citrus Pathology 1, 1–78.

Bové, J. M. & Cassin, J. P. (1968). Problèmes de l’agrumiculture malgache; compterendu de mission. Document IRFA: 52 p.

Doddapaneni, H., Liao, H., Lin, H., Bai, X., Zhao, X., Civerolo, E., Irey, M., Coletta-Filho, H. & Pietersen, G. (2008). Comparative phylogenomics and multi-gene cluster analyses of the Citrus Huanglongbing (HLB)-associated bacterium Candidatus Liberibacter. BMC Res Notes 1, 72.

Garnier, M., Jagoueix-Eveillard, S., Cronje, P., Le Roux, H. & Bové, J. (2000). Genomic characterization of a liberibacter present in an ornamental rutaceous tree, *Calodendrum capense*, in the Western Cape province of South Africa. Proposal of “*Candidatus* Liberibacter africanus subsp. capensis”. Int J Syst Evol Microbiol 50, 2119–2125.

Jagoueix, S., Bové, J. M. & Garnier, M. (1994). The phloem-limited bacterium of greening disease of citrus is a member of the a-subdivision of the Proteobacteria. Int J Syst Bacteriol 44, 379–386.

Jagouiex, S., Bové, J. M. & Garnier, M. (1996). PCR detection of the two “*Candidatus*” liberobacter species associated with greening disease of citrus. Mol Cell Probes 10, 43–50.

Larkin, M. A., Blackshields, G., Brown, N. P., Chenna, R., McGettigan, P. A., McWilliam, H., Valentin, F., Wallace, I. M., Wilm, A. & other authors (2007). Clustal W and Clustal X version 2.0. Bioinformatics 23, 2947–2948.

Lin, H., Doddapaneni, H., Munyaneza, J. E., Civerolo, E. L., Sengoda, V. G., Buchman, J. L. & Stenger, D.C. (2009). Molecular characterization and phylogenetic analysis of 16S rRNA from a new species of “*Candidatus* Liberibacter” associated with Zebra chip disease of potato (*Solanum tuberosum* L.) and the potato psyllid (*Bactericera cockerelli* Sulc). Journal of Plant Pathology 91, 215–219.

Moreno-Enríquez, A., Minero-García, Y., Ramírez-Prado, J. H., Loeza-Kuk, E., Uc-Varguez, A. & Moreno-Valenzuela, O. A. (2014). Comparative analysis of 16S ribosomal RNA of ‘*Candidatus* Liberibacter asiaticus’ associated with Huanglongbing disease of Persian lime and Mexican lime reveals a major haplotype with worldwide distribution. African Journal of Microbiology Research 8, 2861–2873.

Nelson, W. (2012). First record of *Trioza vitreoradiata* (Maskell)(Hemiptera: Triozidae) in citrus. Citrus Research & Technology 33, 35–38.

Nelson, W. R., Fisher, T. W. & Munyaneza, J. E. (2011). Haplotypes of “*Candidatus* Liberibacter solanacearum” suggest long-standing separation. European Journal of Plant Pathology 130, 5–12.

Nelson, W. R., Munyaneza, J. E., McCue, K. F. & Bové, J. M. (2013). The Pangaean origin of “*Candidatus* Liberibacter” species Journal of Plant Pathology 95, 455–461.

Nelson, W. R., Wulff, N. A. & Bové, J. M. (2015). Intragenomic homogeneity on Liberibacter 16S rDNA confirms phylogeny and explains ecological strategy. *bioRxiv* doi: http://dx.doi.org/10.1101/016188.

Nelson, W., Sengoda, V., Alfaro-Fernandez, A., Font, M., Crosslin, J. & Munyaneza, J. (2013). A new haplotype of “*Candidatus* Liberibacter solanacearum” identified in the Mediterranean region. European Journal of Plant Pathology 135, 633–639.

Phahladira, M., Viljoen, R. & Pietersen, G. (2012). Widespread occurrence of “*Candidatus* liberibacter africanus subspecies capensis” in *Calodendrum capense* in South Africa. European Journal of Plant Pathology 134, 39–47.

Pietersen, G., Arrebola, E., Breytenbach, J., Korsten, L., le Roux, H., la Grange, H., Lopes, S., Meyer, J., Pretorius, M. & other authors (2010). A survey for ‘*Candidatus* Liberibacter’ species in South Africa confirms the presence of only ‘*Ca*. L. africanus’ in commercial citrus. Plant Dis 94, 244–249.

Roberts, R., Steenkamp, E. T. & Pietersen, G. (2014). Novel lineages of ‘*Candidatus* Liberibacter africanus’ associated with native Rutaceae hosts of *Trioza erytreae* in South Africa. Int J Syst Evol Microbiol, doi:10.1099/ijs.0.069864-0.

Stackebrandt, E. A. & Goebel, B. M. (1994). Taxonomic note: a place for DNA-DNA reassociation and 16S rRNA sequence analysis in the present species definition in bacteriology. Int J Syst Bacteriol 44, 846–849.

Teresani, G. R., Bertolini, E., Alfaro-Fernandez, A., Martínez, C., Tanaka, F. A. O., Kitajima, E., Rosello, M., Sanjuan, S., Ferrandiz, J. C. & other authors (2014). Association of ‘*Candidatus* Liberibacter solanacearum’ with a vegetative disorder of celery in Spain and development of a real-time PCR method for its detection. Phytopathology 104, 804–811.

Wang, N. & Trivedi, P. (2013). Citrus Huanglongbing: a newly relevant disease presents unprecedented challenges. Phytopathology 103, 652–665.

Weisburg, W., Barns, S., Pelletier, D. & Lane, D. (1991). 16S ribosomal DNA amplification for phylogenetic study. J Bacteriol 173, 697–703.

Woese, C. & Fox, G. (1977). Phylogenetic structure of the prokaryotic domain: The primary kingdoms. Proceedings of the National Academy of Sciences 74, 5088–5090.

